# Action potential propagation speed compensates for traveling distance in the human retina

**DOI:** 10.1101/2024.04.30.591867

**Authors:** Annalisa Bucci, Marc Büttner, Niklas Domdei, Federica Bianca Rosselli, Matej Znidaric, Roland Diggelmann, Martina De Gennaro, Cameron S. Cowan, Wolf Harmening, Andreas Hierlemann, Botond Roska, Felix Franke

## Abstract

Neural information processing requires accurately timed action potentials arriving from presynaptic neurons at the postsynaptic neuron. However, axons of ganglion cells in the human retina feature low axonal conduction speeds and vastly different lengths, which poses a challenge to the brain for constructing a temporally coherent image over the visual field. Combining results from microelectrode array recordings, human behavioral measurements, transmission electron microscopy, and mathematical modelling of the retinal nerve fiber layer, we demonstrate that axonal propagation speeds compensate for variations in axonal length across the human retina including the fovea. The human brain synchronizes the arrival times of action potentials at the optic disc by increasing the diameters of longer axons, which increases their propagation speeds.

## Introduction

To construct a temporally consistent model of the world, the brain needs to integrate information of simultaneous events in the world from sensory modalities with different temporal characteristics, e.g., different signal generation or propagation speeds. Even within a single sensory modality, information from different parts of the sensory space might arrive at different times. The precise relative timing of this information at the integration stage can be highly relevant: In the auditory cortex, the brain can extract behaviorally relevant information about the location of a sound source from the relative timing of arriving action potentials from both ears of less than 1 µs (*1*). When integrating information from both eyes, timing differences below 10 ms are relevant for depth perception (*2, 3*).

The human eye features a diameter of 25 mm, and the retina spans an area of approximately 1,100 mm^2^ (*4*). Locally, the retinal circuitry processes visual signals within each small image patch synchronously across the entire retina, resulting in the generation of action potentials by retinal ganglion cells (RGCs). To travel from the eye to the brain, these action potentials must reach the optic disc, where the optic nerve begins and exits the eyeball. RGCs extend their axons to the brain, but the intraretinal lengths of these axons depend on the specific location of the RGCs within the retina, ranging from a few hundred micrometers near the optic disc to over three centimeters in the periphery (Fig. 1).

**Fig. 1:**
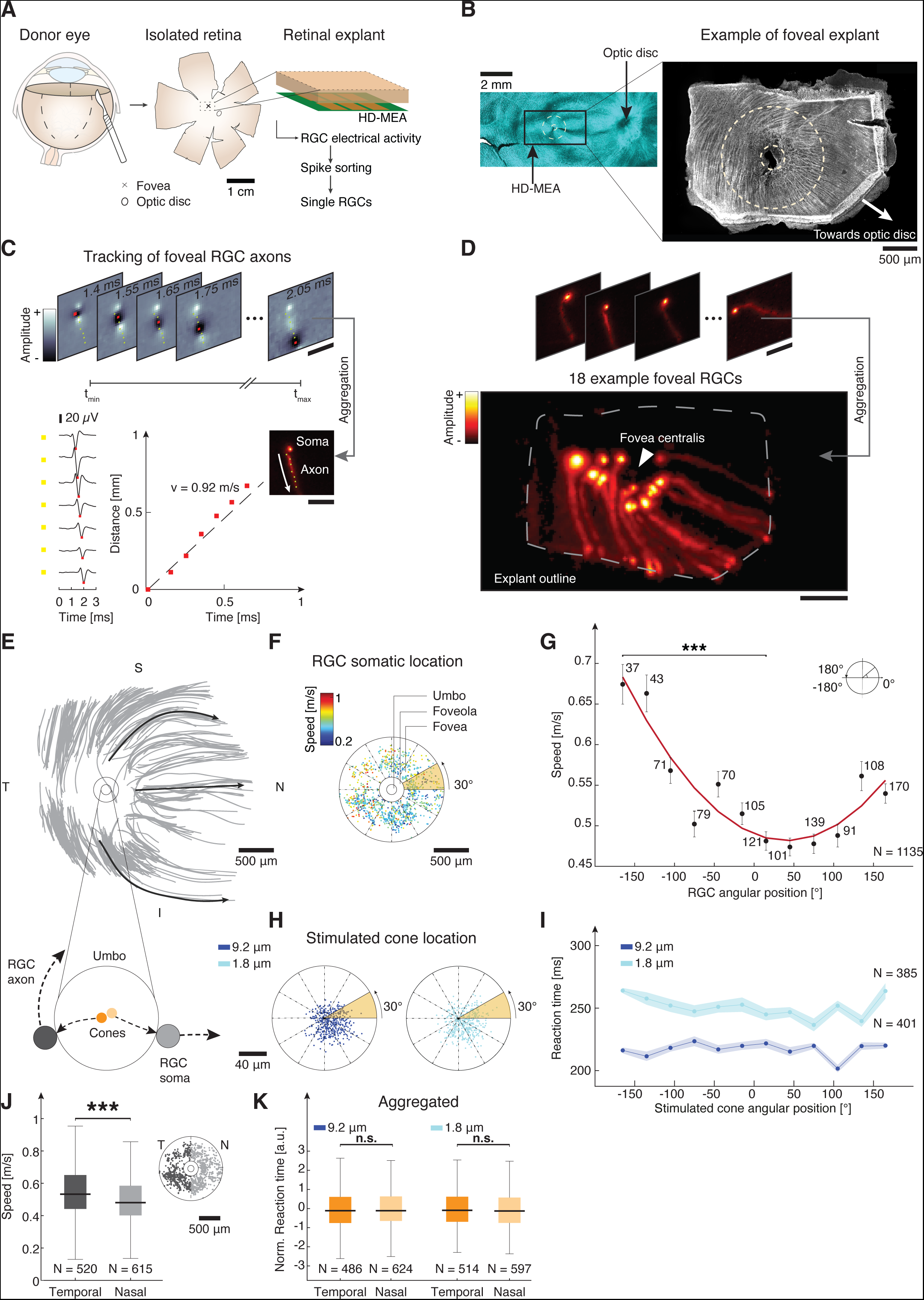
Action potential propagating speeds systematically vary around the human fovea centralis. (**A**) Schematic of sample preparation. Retinal explant is isolated, (cross, fovea; ellipse, optic disc) and positioned onto the HD-MEA. (**B**) Left, immunostaining of flattened axon bundles with Beta III-Tubulin (cyan) in whole-mounted human retina. For comparison, an outline of HD-MEA active array is depicted as a black rectangle (scalebar, 2 mm). Right, immunostained (Beta III-Tubulin) foveal explant after HD-MEA recording. Fovea centralis is located approximately in the center of the explant (scalebar, 500 μm). (**C**) Top sequence: Successive frames of the video depicting the average action potential waveform of an example foveal RGC axon (1.4 ms to 2.05 ms after initiation). Color represents the amplitude of the waveform. Aggregation of individual frames yields an electrical image of the RGC showing the trajectory of the AP (scalebar, 500 µm). Bottom left: Average AP waveform of individual electrodes at increasing distances from the RGC soma (yellow squares, scalebar, 20 µV) for one example RGC. Red square: waveform trough. Bottom right: Linear regression (dashed line) of travel time vs. travel distance for the depicted example. (**D**) Top sequence: Electrical images of representative foveal RGCs. Bottom panel: composite image illustrating the spatial distribution of 18 representative RGCs within the ex vivo retinal explant. White dashed lines delineate the explant outline (scalebar, 500 µm). (**E**) Top: Estimated trajectories of RGC axons for a single preparation. S - superior, N - nasal, I - inferior, T - temporal. Three selected axons are accentuated (black arrows) to illustrate distinct routing patterns. Small circle: Outline of the umbo (magnified below). Large circle: Foveola. Scale bar, 500 µm. Bottom: Schematic of the cone-to-RGC connectivity in the umbo (central black circle). Two adjacent cones (small yellow circles) and their downstream target RGCs (large grey circles) are shown. (**F**) RGC somatic locations (dots) of 1135 RGCs (from N = 11 explants) color coded by their axonal propagation speeds. Radial lines segment the area into 30° angular bins (scalebar, 500 µm). (**G**) Mean axonal propagation speed of RGCs located inside the angular bins depicted in F. Red line: Best fitting sinusoid, added as a visual aid. Pairwise comparisons between each of the 12 bins accounting for multiple comparisons (Kruskal-Wallis test). Significant (***p < 0.001) difference between -165° (0.67 ± 0.02 m/s) and 15° (0.51 ± 0.01 m/s). Error bars indicate ± standard error of the mean (SEM). (**H**) Locations of light stimulation inside the foveola of a healthy human participant using an adaptive optics scanning light ophthalmoscope (AOSLO). Each dot represents the location of the stimulation of a single trial with respect to the cone-density centroid (CDC). Radial lines: 30° angular bins (scale bar, 40 µm). (**I**) Mean reaction times (reported by button press) for the trials shown in H. 9.2 µm, N = 418; 1.5 µm, N = 395. Shaded areas: Mean ± SEM. For each dataset, pairwise comparisons yielded no significant difference between the 12 angular bins (Kruskal-Wallis test; large spot, smallest observed p = 0.72; small spot, largest observed p = 0.99). (**J**) Distribution of axonal propagation speeds for RGCs in the temporal fovea (dark grey, N = 520) and nasal fovea (light grey, N = 615). Inset: Data from F replotted and color coded by spatial bin (temporal vs. nasal). Median speeds differed between temporal (0.53 m/s) and nasal (0.48 m/s) RGCs (Wilcoxon rank sum test, ***p < 0.001). (**K**) Normalized reaction times show no significant median reaction time difference for large (Wilcoxon rank sum test, n.s., p = 0.43) and small (Wilcoxon rank sum test, n.s., p = 0.38) light squares.

Additionally, the region of the retina responsible for high-acuity vision, the fovea centralis, lacks RGCs and their axons (*5, 6*). Consequently, RGC axons must loop around this area, resulting in further increased lengths. In the eye, RGC axons are unmyelinated and exhibit action potential propagation speeds ranging from 0.72 m/s to 1.65 m/s (*7*), which is considerably slower than their speeds observed in the optic nerve, where the RGC axons become myelinated, and conduction speeds can exceed 40 m/s (*8*). The low intraretinal conduction speed, together with the large disparity of axonal lengths, could lead, in the brain, to a significant temporal dispersion of arrival times of originally synchronous action potentials.

Even for RGCs that convey electrical signals of neighboring photoreceptors within the fovea centralis, the axonal length can differ significantly. In the umbo, the very center of the fovea, photoreceptor axons connect radially outwards to displaced bipolar cells within the foveal shoulder, which - in turn - connect to RGCs arranged in a ring-like structure around the umbo (Fig. 1) (*9*). As neighboring photoreceptors can connect to RGCs on opposite sides of this ring, proximity in visual space does not imply proximity in anatomical space (Fig. 1). However, human subjects do not perceive the temporal dispersion of signals from different parts of the visual field, which raises the question of how a compensation for the different travel distances is achieved.

Indirect measurements of axonal conduction speed, based on patterned electroretinograms in humans, indicate a positive correlation between axonal length and conduction speed across different retinal locations (*10*). The axonal diameter is the main factor determining propagation speeds of unmyelinated axons, with larger axons featuring higher propagation speeds (*11, 12*). For human and marmoset retinae, the average axon diameter correlates positively with eccentricity (*13–15*), and foveal RGC axons feature on average smaller diameters than RGCs in the periphery (*16*). This may indicate that axons of RGCs at greater eccentricities, which feature longer axons than RGCs situated closer to the optic disc, also feature higher propagation speeds. However, the two main RGC types in the primate retina, midget and parasol cells (*17, 18*), have different axonal diameters, different axonal propagation speeds, and their relative abundance depends on retinal location (*19*). Parasol cells have larger-diameter axons and conduct significantly faster than midget cells (*7*). Therefore, the relative numbers of the sampled cell types can be a confounding factor when interpreting differences in average axon diameters and propagation speeds at different retinal locations. Furthermore, because RGC axons can take non-straight pathways from soma to optic disc, the intraretinal length of these axons is not a simple function of eccentricity, but strongly depends on retinal location (Fig. 1 B, E).

Mathematical models have been proposed to describe axonal trajectories around the central human retina (*20*). However, the underlying organizational principle of these trajectories remains poorly understood, and precise measurements of axonal lengths have not been conducted for the entire human eye. The temporal dispersion of the arrival times at the optic disc of action potentials that have been elicited synchronously in the retina is an unresolved issue of high relevance for central human vision, as the precise timing of action potentials from presynaptic neurons reaching postsynaptic neurons is crucial for neural information processing (*21, 22*).

## Results

### Axonal propagation speed depends on RGC soma location inside the human fovea

To measure the time necessary for action potentials (APs) of foveal RGCs to reach the optic disc at high spatial and temporal resolution, we recorded the spiking activity of RGCs in the human fovea by means of complementary-metal-oxide-semiconductor (CMOS)-based, planar high-density microelectrode arrays (HD-MEAs) (*23–26*). We dissected donor eyes to isolate the entire retina (Fig. 1A), subsequently resected retinal explants of approximately 3 mm by 2 mm in size, containing the fovea, and placed them - RGC-side down - onto the microelectrode array. The preparation enabled simultaneous recordings of RGC action potentials from foveola, fovea, parafovea and a small portion of the perifovea. After the recordings, we immunolabeled the RGC axon bundles to ascertain the presence of the fovea centralis within the resected explants (Fig. 1B).

We identified the electrical activity of individual neurons through offline spike sorting (*27*) of the electrical recordings and reconstructed the electrical image (average electrical waveforms of APs per electrode) for each neuron across the entire chip (∼26.000 electrodes) at a sampling rate of 20 kHz (Fig. 1C). Superimposing a subset of the electrical images revealed the location of the fovea centralis on the chip and the ring-like arrangement of RGC somas (Fig. 1D). We visualized the electrical images of individual RGCs as videos with a frame rate of 20 kHz. In each video, APs became visible as voltage deflections traveling across the electrodes of the HD-MEA surface (Fig. 1C).

We tracked 1135 individual foveal RGC axons over distances of up to 1.7 mm (10 donors, 11 explants) and calculated the axonal propagation speeds through linear regression of the travelled distance versus travel time for each RGC (Fig. 1E). We registered axonal trajectories of different preparations in a reference coordinate system by aligning the location of the fovea centralis and the orientation between all resected retinal explants. This procedure revealed the axonal wiring-pattern around the fovea centralis, which closely resembled the pattern visible in immunolabelled RGC axon bundles (Fig. 1B). Fig. 1F shows the somatic locations of all tracked foveal RGCs. To quantify the dependence of the axonal AP propagation speed on the RGC soma location within the ring around the fovea centralis, we binned the angular location of the RGC somas in twelve 30° bins (Fig.1F) and compared the average speeds within the bins (Fig. 1G). This approach revealed a strong dependence of the AP propagation speed on the angular location of the foveal RGCs. Specifically, APs of RGCs situated temporal to the umbo (i.e., away from the optic disc) propagated over 40% faster in comparison to RGCs situated nasal to the umbo (i.e., closer to the optic disc).

### Human reaction times to single cone stimulation are homogeneous within the fovea centralis

Within the retina, foveal RGC axons originating in locations temporal to the fovea centralis are significantly longer than those originating on the nasal side and extending directly towards the optic disc (Fig. 1B). We investigated whether the observed increase in AP propagation speed of these axons may compensate for their greater length.

We refer to this as the ‘equal propagation time hypothesis’ suggesting a mechanism that synchronizes AP arrival times at the optic disc for APs initiated simultaneously across the fovea centralis. In contrast, under an ‘equal propagation speed hypothesis’, APs from RGCs with longer axons would feature delays in arrival times at the optic disc, which would potentially increase human reaction times to localized visual stimuli. Previous studies evidenced an increase in human reaction times to localized visual stimulation with greater eccentricity from the fovea (*28*). To test if human reaction times to localized foveal stimulation align with the equal propagation time hypothesis, and to ensure precise and selective stimulation of the densely packed cones within the fovea centralis, we conducted a series of psychophysical experiments using adaptive optics scanning laser ophthalmoscopy (AOSLO) (*29*). We measured the temporal dispersion of human reaction times (RTs) in response to brief flashes of small squares (1.8 µm x 1.8 µm or 9.2 µm x 9.2 µm) of light presented in the umbo.

RTs were quantified by measuring the time interval between onset of the light flash and the pressing of a button by three participants. Responses of one participant are depicted in Fig. 1H-I (mean RT to 1.8 µm squares: 250 ± 42 ms; mean RT to 9.2 µm squares: 218 ± 28 ms), while results aggregated from all participants are presented in Fig. 1K. We used the cone density centroid (CDC), which represents the anatomical location of the center of the foveal pit (*30*), as the center of the fovea (using the preferred retinal locus of fixation (PRL) as center did not change the results qualitatively; data not shown). Similar to our method of sampling the angular positions of RGCs around the fovea centralis, we assessed the angular positions of the stimulation locations in the umbo by grouping them into twelve 30° bins relative to the CDC, as depicted in Fig. 1H. The results showed no significant differences in RTs across the different angular coordinates, although with low statistical power due to the limited number of trials and the inherent variability of RTs. To enhance statistical power, we normalized each participant’s data by subtracting their mean RT and dividing by their respective standard deviation. Subsequently, we grouped the stimulation locations into two regions relative to the CDC: temporal and nasal.

Again, we observed no significant difference, this time with sufficient statistical power to limit the effect size to less than 5.5 ms and 7.5 ms for the large and small squares respectively. This finding aligns with the equal propagation time hypothesis.

### Axonal propagation speeds increase with eccentricity

To understand if axonal AP propagation speeds also vary across different regions of the peripheral retina, which exhibit large disparities in axonal lengths, we measured propagation speeds at different locations and eccentricities. To this end, we measured RGC action potentials across the human and non-human primate (*macaca fascicularis*) retina with the same method as described above but with explants isolated at different retinal locations not including the fovea.

We recorded signals from 16 peripheral human retinal explants, isolated along the naso-temporal axis from 9 donors, which yielded a total of 1186 tracked human peripheral RGC axons (v = 1.13 ± 0.30 m/s, v_max_ = 2.31 m/s, v_min_ = 0.16 m/s, max tracked length: 3.06 mm). Fig. 2B illustrates the average axonal propagation speed along the naso-temporal axis of the human retina. The propagation speed measured in foveal explants was lowest (indicated by the arrow in Fig. 2B, N = 1285) and increased as the eccentricity from the fovea grew. In the far periphery (>10 mm distance from the foveal pit, > 30° eccentricity), the increase in speed was less pronounced. We obtained similar results for macaque retinae evidencing propagation speeds of similar magnitude and dependence of the retinal location. In macaques, we recorded the electrical activity of 128 foveal (4 explants) and 1354 peripheral (8 explants) RGCs along the naso-temporal axis (Fig. 2D). In macaques, we also recorded at four equi-eccentric but radially distant locations (superior, inferior, temporal, nasal) in the far periphery. Although the distributions of propagation speeds featured some differences among the four locations, the average speeds were of similar magnitude (Fig. 2C).

**Fig. 2:**
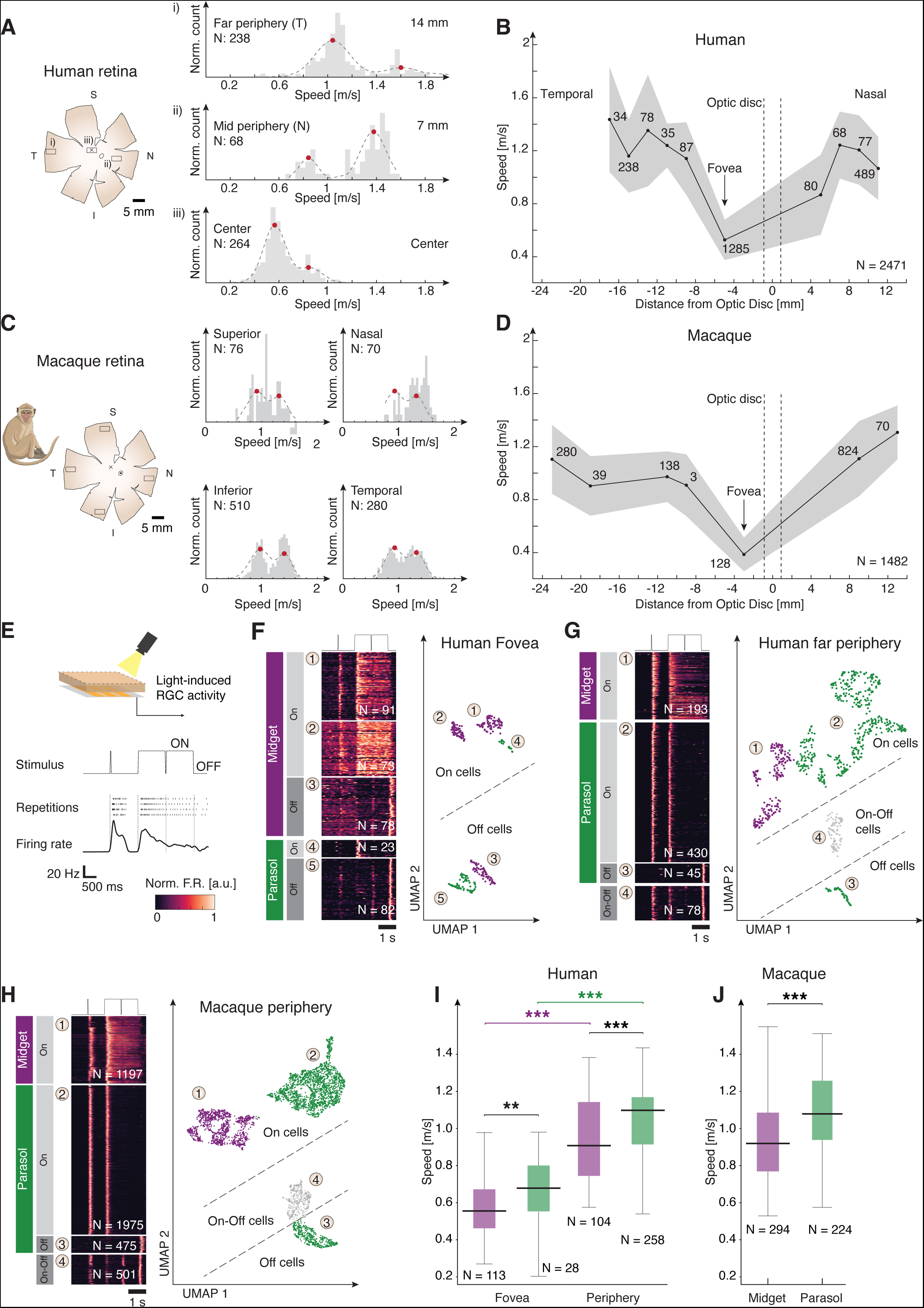
Functional cell typing reveals dependence of axonal propagation speed on eccentricity in midget and parasol cells and in human and macaque retina. (**A**) Histograms of normalized AP speed from three different locations in the human retina (schematic on the left) show bimodal distributions. Dashed line: Fit of Gaussian mixture model with two components; Red dots: Position of peaks. Far periphery (i, peaks: 1.04 m/s and 1.6 m/s, N = 238), mid periphery (ii, peaks: 0.84 m/s and 1.40 m/s, N = 68), and center (iii, peaks: 0.56 m/s and 0.84 m/s, N = 264). (**B**) Mean AP speeds in human retinal explants along the naso-temporal axis by distance from the optic disc; temporal (negative x-axis) vs. nasal (positive x-axis). Numbers indicate the RGC count; shaded region: mean ± SEM. Dashed vertical lines mark optic disc boundaries. (**C**) Histograms of normalized AP speeds in macaque retina (4 explants from the far periphery). S – Superior (peaks: 0.92 m/s and 1.3 m/s), N - nasal (0.93 m/s and 1.3 m/s), I - inferior (0.98 m/s and 1.4 m/s), T - temporal (0.92 m/s and 1.3 m/s, N = 280). Dashed line: Fit of Gaussian mixture model with two components; Red dots: Position of peaks. (**D**) Same as B but for macaque retina. (**E**) From top to bottom: Schematic of HD-MEA recordings with light stimulation. ‘ON/OFF’ light stimulus (contrast over time); Raster plot of spiking response of a representative On midget RGC. Each row represents a single stimulus presentation, each small vertical dash represents a spike; Average firing rate over trials depicted below. (**F-H**) Clustering of light-induced RGC responses to identify functional cell types of the human fovea (F), human periphery (G), and macaque periphery (H). Left: Normalized firing rates (averaged over trials) of all RGCs depicted as rows in response to the stimulus depicted above. Cells were grouped by clusters (number in circle). Labels on the left indicate the putative cell type for groups of clusters. Right: Functional clusters (UMAP projection). UMAP coordinates in F-H were rotated to reflect similarity in cluster structure. Each dot represents an RGC. Colors correspond to cell type (see left). Numbers in circle indicate the cluster number. (**I**) Distributions of AP speeds of midget (purple) and parasol (green) cells in human fovea and periphery. Foveal (midget: 0.56 ± 0.17 m/s; parasol: 0.68 ± 0.17 m/s; median ± SD); Peripheral (midget: 0.91 ± 0.22 m/s; parasol: 1.10 ± 0.20 m/s; median ± SD). Wilcoxon rank sum test: Foveal **p < 0.01, peripheral ***p < 0.001. Intra-type comparison shows lower speeds in the fovea than the periphery. Group medians indicated. (**J**) Same as I but for macaque periphery (median speed midget: 0.9 ± 0.2 m/s; median speed parasol: 1.1 ± 0.2 m/s, Wilcoxon rank sum test, ***p < 0.001).

### Identifying RGC types based on neural responses to light stimulation

The primate retina features two main types of RGCs, midget and parasol cells, which constitute over 90% of all primate RGCs (*17, 18*). Previous work in the macaque retina has shown that peripheral midget cells have lower AP propagation speeds (∼0.8 m/s) compared to parasol cells (∼1.2 m/s) (*31*). Furthermore, the relative number of midget and parasol cells depends on the retinal location (*19*). In the fovea about 90% of RGCs are midget cells, whereas this percentage drops to about 40-45% in the periphery (*32*). Hence, the elevated speed of axonal action potential propagation in peripheral regions may result from the sampling of a greater proportion of parasol cells compared to the fovea.

We investigated whether we could distinguish the two cell type populations in our data. Fig. 2A shows the distributions of propagation speeds measured in three explants originating from three different locations along the naso-temporal axis in the human retina, from the far periphery (∼14 mm from the optic disc), the mid periphery (∼7 mm from the optic disc), and the center (explant centered on the fovea, about 4.7 mm from the optic disc). The distributions were bimodal, and both modes shifted towards lower propagation speeds with decreasing eccentricity. To show that the two distribution peaks indeed corresponded to the two cell types, we measured the light responses of a subset of the recorded RGCs to full-field light stimulation (Fig. 2E). Midget and parasol cells have different roles in primate vision and correspondingly show different response behavior upon stimulation with steps and brief flashes of light, which can be used to identify the cell types (*33–35*). Midget cells show longer sustained responses to steps in the average brightness compared to parasol cells, which produce more transient responses (*33*). We projected a 2-sec-long dark screen, interrupted by a 16-ms-long bright flash, followed by a step to a 2-sec-long bright screen, interrupted by a 16-ms-long dark flash (Fig. 2E). We then represented the neural response of each RGC as a high-dimensional feature vector and used a dimensionality reduction technique to project the high-dimensional data set onto two dimensions (UMAP (*36*), Fig. 2F-H).

Additionally to and independently of the dimensionality reduction, we clustered the RGC data to identify groups of RGCs featuring similar light-evoked responses, and which likely belong to the same cell type. We performed this analysis independently for the three different data sets from human fovea (Fig. 2F), human periphery (Fig. 2G), and macaque periphery (Fig. 2H). We then labelled the groups as midget or parasol cells based on the similarity of the average response within each group and the known response behavior of midget and parasol cells. This way, we classified in total 5241 RGCs. In the three data sets, we could identify response behaviors that can be expected from the main primate RGC types: Cells showing increased activity upon positive contrast changes (On cells), negative contrast changes (Off cells), and cells, which responded to both changes (On-Off cells). The On-Off cell cluster was absent in recordings from the fovea centralis. This finding aligns with prior results indicating that small bistratified cells, characterized by On-Off response behavior, are less prevalent in the fovea (*32*). Additionally, we identified cells exhibiting transient responses to contrast changes, a characteristic trait of parasol cells, as well as cells displaying sustained activity in response to such contrast changes, a typical behavior observed in midget cells (32, 33).

### Midget and parasol cell axons show increased propagation speeds with eccentricity

We measured both the propagation speeds and light responses of 1021 RGCs (human fovea: 141, human periphery: 362, macaque periphery: 518). For these cells, we analyzed the AP propagation speed as a function of cell type (Fig. 2I, J). Across both cell types, axonal AP propagation speeds were higher in the periphery compared to the fovea, and axons of midget cells propagated APs at lower speeds than axons of parasol cells. Specifically, in the human foveal region, midget and parasol cells featured median propagation speeds of 0.6 ± 0.2 m/s and 0.7 ± 0.2 m/s, respectively. In both human and macaque periphery, midget cells exhibited median axonal AP propagation speeds of 0.9 ± 0.2 m/s - slower than parasol cells - which showed speeds of 1.1 ± 0.2 m/s. Midget cells in the periphery demonstrated higher axonal AP propagation speeds than parasol cells in the fovea, underscoring the complex interplay between cell types and retinal locations in determining axonal speed. Within a single retinal location, we found a strong association between functional cell type and AP propagation speed, suggesting that AP propagation speed alone is a good indicator to distinguish midget from parasol cells. However, for a reliable speed-based classification, it is necessary to record from RGCs at the same retinal location, as different locations feature vastly different speed distributions, which would then confound the classification.

### A simple model explains the observed axonal trajectories across the entire human retina

So far, we measured axonal propagation speeds as a function of the RGC somatic location and cell type. However, to understand to what degree the observed speed difference compensates for different axonal length, we needed to correlate the measured speed with the intraretinal axonal length. To this end, we developed a mathematical model that described the precise trajectories of all RGC axons across the entire human retina. As a starting point, we used the observation that the pattern of axonal trajectories around the human fovea that is visible in our whole-mount images (Fig. 1B, 3A) resembled field lines of magnetic fields, electrical fields, or streamlines of fluid flow under a laminar-flow regime (Fig. 3C, top left). The field lines are solutions to Laplace’s equation, which is a 2^nd^ order partial differential equation (PDE). Laplace’s equation describes many physical phenomena, including non-turbulent fluid flow, electromagnetic effects, and diffusion. Under steady-state conditions, the local concentration of a diffusing chemical does not change, and consequently, the amount of the chemical that enters a certain spatial compartment must be exactly equal to the amount that leaves this compartment: Δ*c* =0, where *c* is the concentration and Δ the Laplace operator or spatial derivative. When axons grow, they establish their trajectories by following gradients of specific chemicals with their growth cones (*37*). These chemicals are often distributed by diffusion, so that it is plausible to also use Laplace’s equation, which describes diffusion processes, to describe axonal trajectories. Laplace’s equation is linear and, therefore, abides by the superposition principle. Fig. 3C, top illustrates the superposition of a sink and a source (both solutions to Laplace’s equation), which yields a dipole (a third solution). If we modify this example by making the sink stronger than the source, the resulting pattern of field lines changes, and strongly resembles the axonal trajectories around the human fovea. Motivated by this observation, we developed a 3D model of the geometry of the human eye and solved Laplace’s equation for the semi-spherical geometry of the human retina. We placed a weak, but spatially extended source at the location of the fovea, a stronger sink at the location of the optic disc, and another circular source at the rim of the retina (i.e., at the ora serrata) (Fig. 3D). Apart from the geometry, only five parameters specified the entire model: the relative strengths of the two sources and the sink, the spatial extent of the foveal source, and the diffusivity of the retinal tissue. Our model yielded a concentration gradient of a chemical, created at the fovea and ora serrata, and absorbed at the optic disc (Fig. 3D, top right). If an RGC growth cone started at any location in the retina and followed this gradient, it would reach the optic disc along a trajectory determined by the field lines. Thus, the resulting trajectory was the corresponding field line. To test if our model accurately described axonal trajectories in the human retina, we estimated the trajectories of axonal bundles across the human retina in immunolabeled whole-mount retinal images (Fig. 3A, B) with an automated procedure (Fig. 3B). We then fitted the five parameters of our model to the extracted trajectories in the central area containing fovea and optic disc and compared the model to the data (Fig. 3E, F, G). Despite the low number of parameters, the model described the axonal trajectories qualitatively and quantitatively well (Fovea 1: R^2^ = 0.91; Fovea 2: R^2^ = 0.95).

**Fig. 3:**
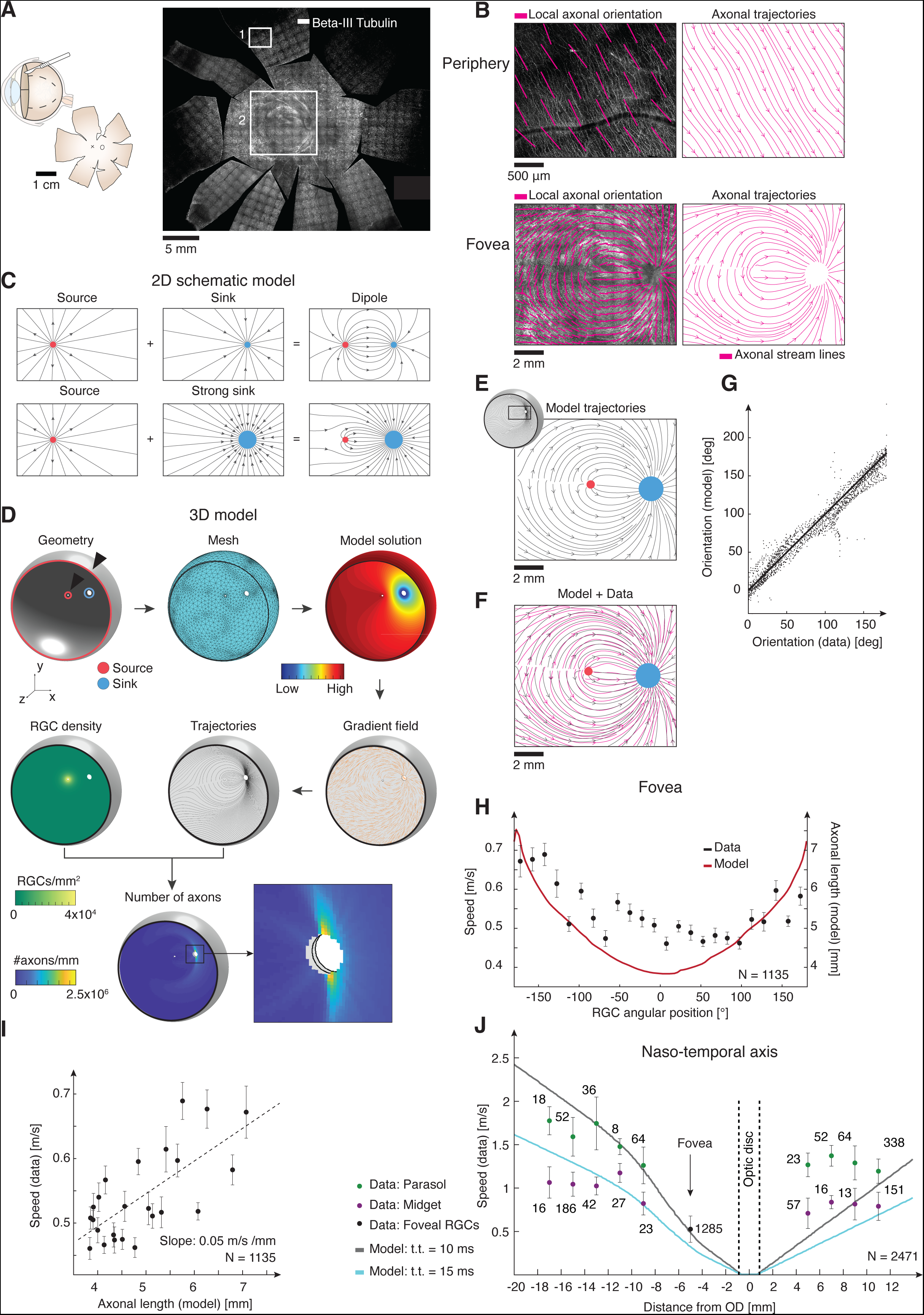
A model based on Laplace’s equations demonstrates that axonal propagation speed correlates to axonal length. (**A**) Schematic of tissue preparation (left) and human whole-mount retina immunostained for axon bundles with Beta-III tubulin (right). Two white rectangles depict the regions highlighted in (D-E). (**B**) Top, estimation of axonal trajectories from the immunostained whole-mount shown in (A). Left, zoom into the image in (A) at a peripheral location superimposed with the estimated local axonal orientation (short magenta line segments). Right, estimated axonal trajectories (magenta lines) at that location. Bottom, same as top, but for a region centered on the fovea. (**C**) Examples of solutions to Laplace’s equations and how they linearly combine to yield new solutions. Black lines indicate field lines of the underlying potential, i.e., trajectories a particle would follow if it was transported along the gradient of the potential. Red dot: Source; Blue dot: Sink. (**D**) Schematic of the modelling process for the three-dimensional model of the axonal trajectories in the human retina. Top row: (i) Specification of the geometry of the eye, including location and size of sources (red circles, highlighted by black triangles) and sink (blue circle); (ii) Generated mesh for numerical solution of Laplace’s equation. (iii) The solution is a scalar field representing the concentration of a chemical that is produced at the ora serrata and at the fovea, diffusing across the retina, and is absorbed at the optic disc. Middle row (right to left): (iv) Orientation of the resulting concentration gradient, guiding axonal growth, depicted as short orange line segments; (v) Example axonal trajectories that follow the concentration gradient (black lines); (vi) Density of RGCs across the retina used to calculate the number of axons that cross each location of the retina. Bottom left: Number of axons at each location within the retina. Inset shows a zoom on the region containing the optic disc (small black square). (**E**) Zoom on the foveal region of the modeled axonal trajectories corresponding to the foveal region shown in B (bottom). Black lines indicate modelled axonal trajectories. (**F**) Superposition of the modeled trajectories shown in E and of the trajectories estimated from the whole-mount shown in (B, bottom right). (**G**) Comparison of modeled trajectories to estimated trajectories (data) for the region shown in F. Solid line indicates unity. Each point represents the local orientations in a small image patch. (**H**) Foveal speed data presented in Fig. 1G binned every 15° (data reported as mean ± SEM, black, axis on the left) overlaid with model axonal length (solid red line, axis on the right). (**I**) Same speed data reported in (H) reported as mean ± SEM and plotted against model axonal. Dashed line, linear regression fit. (**J**) Data replotted from Fig. 2B. Each data point (except for foveal RGCs, black) is assigned to midget (purple) and parasol (green) cell populations. Numbers indicate the number of RGCs per bin; data reported as mean ± SEM. Dashed vertical lines mark optic disc’s boundaries. Solid lines represent speed corresponding to 100% compensation for a travel time (t.t.) of 10 ms (grey) and 15 ms (cyan).

We verified the model’s validity by predicting the thickness of the retinal nerve fiber layer (RNFL). The RNFL is the innermost retinal layer, consisting of the unmyelinated RGC axons. The more axons pass through a location in the retina, the thicker is the RNFL, and the RNFL thickness can be assessed in vivo by optical coherence tomography (OCT) (*38*). We modeled the RGC density as a function of retinal location based on measurements of primate RGC densities (*39, 40*). We arranged the axons along the field lines of our 3D model, determined the respective axon densities, and counted, for each location within the RNFL, how many RGC axons passed through that location. The resulting axon densities agreed qualitatively with measurements of the RNFL thickness in healthy subjects (Fig. S2B).

### Axonal propagation speed compensates for intraretinal axonal length

We then used the model to correlate axonal AP propagation speeds with intraretinal axonal length with the aim to estimate the intraretinal travel time of APs from RGC somas to the optic disc. In the following, ‘axonal length’ refers to the intraretinal axonal length as defined by the model.

In the ring-like structure around the fovea centralis (radius 0.25 mm), axonal lengths ranged from a minimum of 3.8 mm on the temporal side to a maximum of 7.8 mm on the nasal side (Fig. 3K). These values demonstrate that under the ‘equal speed hypothesis’ (where all APs travel at the same speed), APs starting on the temporal side of the fovea centralis would take twice as long to reach the optic disc compared to those starting on the nasal side.

At a speed of 0.4 m/s, this would result in 19.5 ms travel time for temporally located RGCs vs. 9.5 ms for nasally located RGCs. However, the measured AP propagation speeds correlated with the modelled axonal lengths (Fig. 3J, K) so that the difference between minimal and maximal travel times (‘temporal dispersion’) amounted to less than 2.5 ms. A correlation between propagation speed and eccentricity also existed in the periphery for midget and parasol cells (Fig. 3L); i.e., longer axons showed higher propagation speeds.

### Axonal thickness determines axonal propagation speed

A main factor determining axonal propagation speed in unmyelinated axons is their thickness or diameter, with larger diameters reducing axial resistance and thereby enhancing conduction speeds. This relationship scales proportionally with the square root of the axon diameter (*12*). We explored the role of thickness in unmyelinated RGC axons in relation to the speed of AP propagation. Previous findings have highlighted a positive correlation between retinal eccentricity and axonal diameter, with axons of more eccentric RGCs being thicker (*16*). Notably, peripheral primate parasol cells exhibit higher propagation speeds (*41*) and thicker axons than midget cells (*42*). To assess if variations in axonal thickness could account for the observed differences in AP propagation speed around the human fovea centralis, we employed Transmission Electron Microscopy (TEM) to measure RGC axonal diameters at four different retinal locations around the fovea centralis. We first isolated explants (∼1 mm^2^ in size) from the central region of postmortem human retinae as shown in Fig. 4A. We then processed these samples into 70 nm thick cross-sections of resin-embedded tissue, which were subsequently imaged with TEM at 8500x magnification. Each image covered patches of RNFL ranging from 100 to 200 µm in size. Post-imaging, we segmented the images to distinguish the RGC axon cross-sections and delineate their outlines (Fig. 4B, top). Diameter estimates were derived by fitting ellipses to these outlines and measuring their minor axes (Fig. 4B, bottom), which resulted in over 110k human retinal axon diameters. This analysis revealed bimodal diameter distributions at the four different locations, which we fitted utilizing Gaussian mixture models with two components (Fig. 4C). The first component represented axons of small diameter and high numerical abundance, whereas the second component represented axons with larger diameters, small abundance, and larger variability in diameter. For each of the four locations, we calculated the corresponding axonal length utilizing our model, and we then related the model axon length to the average axon diameters (Fig. 4 D, E). For both components, axon diameters positively correlated with intraretinal axon lengths, and the average diameter of axons increased by over 70% when comparing lengths of 4.7 mm to 8 mm (Fig. 4D). These results were in qualitative agreement with our AP propagation speed measurements (Fig. 3I).

**Fig. 4:**
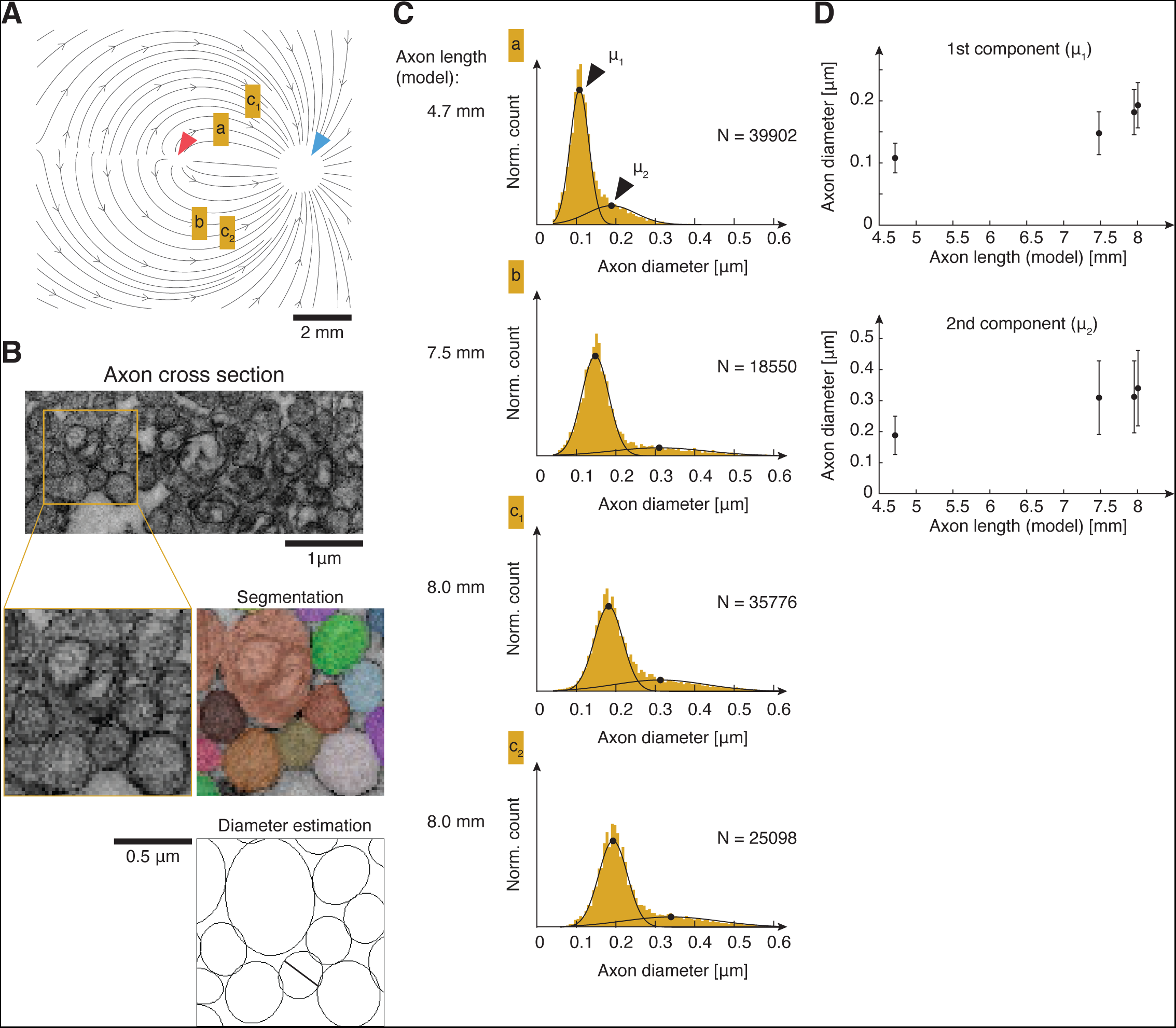
RGC Axon diameters increase with intraretinal axonal length. (**A**) RGC axon sampling locations (a, b, c_1_, c_2_) overlaid with model axonal trajectories. (Red triangle: Fovea. Blue triangle: Optic disc. Scale bar, 2 mm). (**B**) Top, Example of TEM image of RNFL axons in cross-section. Below: Zoom into the TEM image (left) and corresponding segmentation (right, colors indicate different axons). Bottom: Outlines of segmented axons. Black line indicates the minor axis of an ellipse fit to one axon outline (Scalebar, 1 µm). (**C**) Histograms of RGC axon diameters for four different locations sampled from one central human retina. Model axonal length for the four locations marked on the left of each histogram. Numbers indicate the number of estimated RGC axon diameters per location. Black lines: Fit of Gaussian mixture model with two components. Black dots: Mean value of each component (μ1 and μ2). (**D**) Mean ± SD of the two components fit in C versus model axonal length.

## Discussion

By a combination of a set of experimental approaches with a model of the RNFL, we could relate axonal propagation speed, intraocular length, and diameter with RGC soma location, and functional RGC type across the human retina. We showed that in human and macaque, intraretinal axonal length positively correlated with axonal diameters and axonal propagation speeds. This correlation reduced the temporal dispersion of APs at the optic disc (i.e., the difference between the arrival times of two APs elicited by simultaneous visual events) and thus compensated for the variation in axonal lengths of different RGCs. For APs initiated in the human fovea, this compensation reduced the temporal dispersion at the optic disc to below 2.5 ms, which was consistent with our behavioral measurements of human reaction times.

We found a similar correlation between length and propagation speed in the periphery, although for larger eccentricities the speed did not fully compensate the increased axonal lengths (Fig. 3J) and could be a cause for the previously observed increased human reaction times for visual stimulation in the periphery (*28*).

Our findings highlight the importance of relative axonal propagation delays for precise AP timing even in unmyelinated axons within the human brain. Precise relative delays of APs from different neurons can be highly relevant for perception, e.g., in the auditory system, where sub-millisecond timing among APs from different neurons enables precise sound localization (*1*). Our work supports the hypothesis that the human brain modulates axonal propagation speeds to enhance neural information processing.

## Acknowledgments

We would like to acknowledge the following individuals and organizations for their valuable contributions to our research: We extend our heartfelt appreciation to the organ and tissue donors, as well as their families, whose benevolent contributions have greatly advanced scientific knowledge. We are grateful to the transplant coordinators of the Universitätsspital Basel for their dedicated support in facilitating the organ and tissue donation process. We acknowledge Cinzia Tiberi Schmidt from Bio-EM for expertise in electron microscopy sample preparation and image acquisition; Livia Friedlin from the Augenklinik for providing post-mortem tissue; SILABE (Simian Laboratory Europe, University of Strasbourg) for their collaboration and provision of primate tissue for our research investigations; Akos Kusnyerik for coordinating the tissue donation process, Mohammad Khani and Helene Schreyer for help with the preparation of the human tissue, and Arjun Bharioke for comments on a draft of the manuscript.

This study was financially supported by the Swiss National Science Foundation (SNSF) under Eccellenza Grant (PCEFP3_187001, F.F.), the Projects in Life Sciences Grant (310030_220209, F.F.), Sinergia Grants (CRSII5_173728 and CRSII5_216632; A.B., R.D., M.Z., A.H., B.R.); by the European Commission under the ERC Advanced Grant 694829 (“neuroXscales”, A.H.); by the German Research Foundation under Emmy Noether-Program (Ha5323/5-1, W.H.), Priority Program SPP2127 (Ha5323/6-1, W.H.); and by the Carl Zeiss Foundation under HC-AOSLO (W.H.).

## Author contributions

Conceptualization: FF, AB

Methodology: AB, FF, MB, ND, WH, FBR, MZ, RD, MSG, CSC

Investigation: AB, FF, MB, ND, WH

Visualization: FF, AB

Funding acquisition: FF, AH, BR, WH

Project administration: FF

Supervision: FF, AH, BR, WH

Writing – original draft: AB, FF

Writing – review & editing: AB, FF, AH, ND, WH

## Materials and Methods

### Human retinal tissue

Human eyes were obtained from multi-organ donors with no documented history of eye diseases. Donors, encompassing both sexes, ranged in age from 30 to 80 years. Enucleations were performed by the Augenklinik Basel in collaboration with the University Hospital of Basel. All samples were anonymized. These procedures complied with the Declaration of Helsinki’ principles and were approved by the local ethics committee (‘Ethikkommission Nordwest-und Zentralschweiz EKNZ’).

### Human retinal explant preparation and electrophysiological setup

Post enucleation, the corneal tissue was excised for transplantation purposes, and the vitreous humor was carefully removed following radial incisions on the eye bulbs. Critically, we minimized the time between clamping of the eye’s central artery, which interrupted blood supply, and the subsequent immersion of the eyes in pre-oxygenated (95% O_2_ and 5% CO_2_) Ames’ medium (A4034, Sigma-Aldrich Chemie GmbH, Buchs, Switzerland). The enucleated eyes, or more specifically, the eyecups, were rapidly transported to our laboratory, maintained in an actively oxygenated environment (Minican, art. 800002225, PanGas AG HiQ®, Dagmersellen, Switzerland), consistently under 20 minutes. This rapid processing was critical for maintaining tissue viability for subsequent electrophysiological recordings. Retinal explants were then isolated and flattened via relaxing cuts under dim red-light conditions in oxygenated Ames’ medium at room temperature. Explants, approximately 6 mm^2^ in size, were placed flat on the CMOS high-density microelectrode arrays (HD-MEAs, (*23*)) with the RGC layer facing the electrodes. To enhance signal-to-noise ratio, the explants were affixed to the electrodes using a transparent cell culture membrane (Corning Incorporated, 3450 - Clear), pressed against the photoreceptor layer. To ensure precision and to avoid damaging the retinal circuitry, the membrane was lowered under constant visual inspection using a micromanipulator (MBT616D/M, Thorlabs, New Jersey, United States) connected to a custom device for maintaining the membrane flat. For electrophysiological recordings, we maintained the explants in a constantly perfused oxygenated Ames’ medium and temperature-controlled environment. The Ames’ medium was warmed to 37 °C by a temperature controller (TC01/02, Multi Channel Systems MCS GmbH, Reutlingen, Germany) and delivered at a flow rate of 6 ml/min using a peristaltic pump (BT100-1L, Darwin Microfluidics, Paris, France). Used medium was removed by suction through a centralized vacuum line connected to a syphon system. This setup ensured the viability of the retinal tissue, allowing for extended recording durations of up to 20 hours.

### Non-human primate retinal explant preparation

We utilized retinal explants of 15 healthy adult Cynomolgus macaques (*Macaca fascicularis*). These animals were housed and monitored at the Simian Laboratory Europe (SILABE), Strasbourg, in compliance with the European Directive 2010/63/EU. Retinal explants were sourced from macaques, courtesy of our collaborators. All animals were euthanized for different research projects, which involved treatment of some of the eyes via subretinal injection but did not make use of the complete retinal tissue. All procedures performed on the animals were approved by the Comité Régional d’Ethique en Matière d’Expérimentation Animale de Strasbourg and registered with the following numbers APAFIS#5716_2016061714424948_v6 (2018/08/28), APAFIS#32591_2021072914362019_v5 (2022/04/03), and APAFIS#27357-2020092811266511_v2 (2020/12/28). The enucleation process was conducted under deep terminal anesthesia with ongoing monitoring. It is imperative to perform enucleations prior to euthanasia to maximally preserve vascularized tissues and prevent cellular damage due to oxygen deprivation. The anesthesia and analgesia protocols guaranteed that the animals remained unconscious and free from pain throughout the entire procedure, up until the point of euthanasia. The protocol for enucleation encompassed the following steps: on the night prior to the procedure, the animals were fasted. Subsequently, they were sedated using Ketamine (Ketamine 1000, 10 mg/kg, intramuscularly) and transported to the preparation room. A venous catheter was inserted into the saphenous vein, followed by an intravenous injection of Propofol (Propovet, 5-10 mg/kg) through the catheter. The animals were then intubated and administered isoflurane gas anesthesia (ISOVET, 1-2.5%, inhalation) alongside a potent analgesic, Morphine (MORPHINE AGUETTANT, 2 mg/kg, intramuscularly). After conducting an ocular examination via OCT imaging (Atlantis OCT, Topcon) to ensure the integrity of the eyes, the animals were prepared for enucleation. Local anesthesia was achieved using a procaine-based solution (PROCAMIDOR, 17.3 mg/ml, 0.1 ml/eye, subcutaneously) delivered through 3-4 subcutaneous injections around the orbital area. After enucleation, the animals were euthanized using a lethal dose of Pentobarbital (DOLETHAL; 180 mg/kg, intravenously). Similarly to the preparation of human eyes, the anterior segment and vitreous body were removed immediately after enucleation, resulting in the preservation of the eyecup. Foveal retinal explants were obtained from macaque eyes that had not been treated. Peripheral retinal explants were sourced from macaques that had undergone subretinal injections, which, upon injection, caused the temporary formation of localized blebs. Regions impacted by these blebs were identified and annotated. Our collaborators provided peripheral, untreated areas of the retina for our use (excised with a 4 mm punch, Kai Medical, BPP-40F), chosen to avoid the bleb-affected zones. HD-MEA recordings were performed at two laboratories. A set of experiments was conducted directly on-site at SILABE, Mittelhausbergen, whereas another set required the transport of samples from Mittelhausbergen to Basel. The eye cups destined for Basel were submerged in pre-oxygenated (95% O_2_ and 5% CO_2_) Ames’ medium (A4034, Sigma-Aldrich Chemie GmbH, Buchs, Switzerland) and airlifted by helicopter (Helitrans AG, Basel) to minimize the transit time. Experiments conducted on site did not involve any sample transportation. Under both on-site and transport conditions, akin to the handling of human retinal tissue, isolated macaque retinal explants, each measuring approximately 6 mm^2^, were positioned flat on the HD-MEA for recordings. Throughout our analysis, data from both transport and non-transport conditions were processed in the same manner. Our findings revealed no differences between the two groups; therefore, they were pooled for all analyses.

### Electrophysiological recording using HD-MEAs

We employed CMOS high-density microelectrode arrays (*23*) for the electrophysiological recordings of retinal ganglion cells (RGCs) in ex vivo explants of human and non-human primate retina. These arrays featured a recording area of 3.85 x 2.1 mm^2^ with 26,400 electrodes, spaced at a pitch of 17.5 μm. Signal was acquired via 1,024 recording channels at a sampling rate of 20 kHz.

To assign the extracellular action potentials to individual neurons, we used the off-line automatic spike sorter algorithm as outlined by (*27*). Briefly, electrodes recording electrical activity were grouped into local electrode groups, each comprising up to nine electrodes. The following steps were performed for each group independently and in parallel. The electrical signal from each electrode was bandpass filtered between 0.3 and 6 kHz. Spike detection occurred when the signal surpassed a predefined threshold, set at 4.2 times the standard deviation of the noise level. For each spike, the spatiotemporal waveform was extracted and saved. Spike templates corresponding to different neurons were identified through unsupervised data dimensionality reduction followed by a mean-shift clustering algorithm. Spikes were then matched to the most similar template. Since a neuron could be detected on multiple local electrode groups, duplicate neurons were detected and removed based on the similarity of their average spike waveforms and the timing of their spikes.

### Recording spontaneous spiking activity with HD-MEAs

The HD-MEAs can record from a nearly arbitrary set of 1,024 out of 26,400 electrodes simultaneously. To record spiking activity on all electrodes, we split the recording in different periods, each with a different set of electrodes (‘configurations’). We included a small subset of 45 shared electrodes, which were contained in each electrode configuration. For the first configuration, the remainder of the electrodes was chosen randomly, and then, in each subsequent configuration, the remainder of electrodes was chosen randomly from the set of electrodes not yet included in any electrode configuration. We repeated this process across 29 different configurations. For each configuration, electrical signals were captured for 30 seconds, amounting to a total recording duration of approximately 16 minutes. This strategy allowed us to eventually capture data from the entire array (Fig. S1). Importantly, the subset of electrodes consistently included in every configuration provided a continuous recording across all configurations allowing off-line spike sorting. For each resulting spike sorted RGC, we calculated the average action potential waveform across the entire array by averaging the waveforms on each electrode within each configuration. For each retinal explant, this recording protocol was repeated multiple times, strategically selecting the set of shared electrodes from various regions of interest in the preparation, such as the rim of the fovea centralis and at different eccentricities within the recorded explant.

### Recording light-evoked spiking activity with HD-MEAs

Light stimuli, consisting of full-field contrast steps, were generated, and controlled using Psychtoolbox in MATLAB (*43*). These light stimuli were projected onto the retina using a DLP LightCrafter 4710 projector (Texas Instruments, Dallas, TX, USA) from which the magnifying optics had been removed. The light was focused onto the retina using a Nikon camera lens and a 2.5x customized objective (Thorlabs, Newton, NJ, USA), illuminating an area of 2.5 x 1.9 mm^2^. Specific regions of interest on the retina were identified for recording RGC light-induced spiking activity. Configurations of up to 1,024 electrodes, centered on these regions, were selected for targeted recording. A full-field contrast step stimulus including contrast flashes was used as a light stimulus. The stimulus consisted of the following ‘steps’ and ‘flashes’: (i) 1 s of black, (ii) a single frame (1/60 s) of white (‘flash’), (iii), 1 s of black (iv) then a ‘step’ to 1s of white, (v) a single frame (1/60 s) of black (‘flash’), (vi) 1 s of white, (vii) 0.5 s of black. The stimulus was repeated four times (trials).

### Immunohistochemistry

Post recording, the retinal explant was removed from the HD-MEA chip and was immersed in 4% PFA (paraformaldehyde) for 30 minutes at room temperature and washed overnight in PBS (Phosphate Buffered Saline). The sample was then immersed in 30% sucrose solution (in PBS) for 2 hours, and a cycle of freezing and thawing was repeated 3 times. Samples were afterwards immersed in blocking solution (Table 1) for 2 hours under shaking conditions at room temperature. Subsequently, samples were immersed in primary antibody solution (Table 2 and 3) for 5 days under shaking conditions at room temperature. Samples were washed in PBS 3 times. Secondary antibody solution (Table 2 and 3) was applied for 2 hours under shaking conditions at room temperature. Samples, after being washed in PBS for 3 times, were finally mounted on coverslips by using a glycerol based liquid mountant (ProLong™ Diamond Antifade Mountant, Thermofisher) applied directly on fluorescently labeled tissue (Fig. 1B).

**Table 1:** Blocking solution.

| Reagents | Suppliers | Concentrations |
| --- | --- | --- |
| NDS | Sigma-Aldrich, S30-M | 10% |
| BSA | Sigma-Aldrich | 1% |
| PBS 10x | Gibco | 1x |
| NaN<sub>3</sub> 2% | Sigma-Aldrich, S2002 | 0.02% |
| Triton X 10% | Sigma-Aldrich, 93443 | 0.5% |

**Table 2:** Primary/secondary antibody solutions.

| Reagents | Suppliers | Concentrations |
| --- | --- | --- |
| NDS | Sigma-Aldrich, S30-M | 3% |
| BSA | Sigma-Aldrich | 1% |
| PBS 10x | Gibco | 1x |
| NaN<sub>3</sub> 2% | Sigma-Aldrich, S2002 | 0.02% |
| Triton X 10% | Sigma-Aldrich, 93443 | 0.5% |

**Table 3:** Antibodies.

|  | Primary |  | Secondary |
| --- | --- | --- | --- |
| RNFL | mouse anti-Beta III-tubulin | Millipore, MAB1637, 1:200 | donkey anti-mouse IgG conjugated with Alexa-647, 1:200 |

For immunostaining whole-mount human retinae, entire eye bulbs were preserved in 4% PFA for a minimum duration of five days. This preparatory step ensured the tissues were adequately fixed for the ensuing histological analyses. Following fixation, the specimens underwent a thorough rinsing process in PBS to remove residual fixative. The retinae were then dissected from the surrounding ocular tissues. Relaxation cuts were made to enable the retinae to be laid flat. Given their size, the retinae were sectioned into multiple fragments, each of which was mounted and imaged independently (Fig. 1B, Fig. 3A).

### Confocal microscopy

Images were captured using a Yokogawa spinning disk confocal system attached to an Olympus microscope, operated with CellSens Software by Olympus. A composite image illustrating the human retinal nerve fiber layer (RNFL) shown in Fig. 1B was created by stitching together images taken with a 4x objective lens. For the assembly of Fig. 3A, individual segments of the retina were imaged separately utilizing a 10x objective lens. The processing of these images was performed using ImageJ software (Fiji distribution), and they were seamlessly integrated into a singular image using Adobe Photoshop.

### Analysis of light responses

We conducted the analysis of neural responses to light stimulation using Python 3.8 (libraries included numpy, pandas, scipy and the electrophysiology package elephant (*44*)). Time-dependent firing rates,*r*(*t*), in response to each repetition of a light stimulus (‘trial’) were determined using kernel density estimation (*45*) with a Gaussian kernel (*46*) (*Δt* =10 *ms, σ* =50 *ms*). The firing rates were averaged over trials and normalized by the maximum firing rate for each neuron. To assess each neuron’s responsiveness to light, we assigned a quality index (QI, (*47*)) calculated as

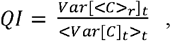

where the indices *r* and *t* indicated taking the expectation or calculating the variance over trials or time bins, respectively. The QI estimated the variability of the neuron’s firing rate across trials relative to the variability of the trial-averaged firing rate. Here, *C* was a *T×R* matrix where T was the number of time bins and R was the number of trials. A QI of 1 indicated that the neuron’s mean response consistently reflected individual trial responses and tended towards 1/*R* when responses over different trials varied significantly. Neurons with a QI lower than 0.45 were omitted from further analysis.

### Clustering

Light-responsive neurons recorded in human fovea (N = 711; 5 retinae), human periphery (N = 1364, 1 retina), and macaque periphery (N = 9385, 7 retinae) were clustered separately (Fig. 2F-H) based on their light-evoked firing rates. The dimensionality of the normalized firing rate vectors corresponded to the number of time bins (T = 450 time bins). Prior to clustering, to reduce the dimensionality, we employed the nonlinear dimensionality reduction technique known as Uniform Manifold Approximation and Projection (UMAP) (*36*) with n_*neighbors* = 10, *min_dist* = 0, metric = ‘Euclidean’, n_components_ = 2. This resulted in two-dimensional feature vectors, each representing the mean response of each light-responsive neuron, which we could visualize in 2D UMAP coordinates. We then performed hierarchical clustering on the two-dimensional feature vectors using the *AgglomerativeClustering* function of the Python package *SKlearn* (*48*) (metric = ‘Euclidean’, linkage = ‘average’). To ascertain the number of clusters, we adopted the approach delineated in (*49*). To determine the optimal clustering, we plotted the number of clusters against the hierarchical clustering algorithm’s merging steps, setting dataset-specific minimum element thresholds per cluster, contingent upon the size of each distinct dataset. We set a minimum of 20 cells per cluster for the human fovea and periphery, a minimum of 100 cells per cluster for the macaque periphery. We stopped the hierarchical clustering algorithm at the merging step that produced the maximal number of clusters that fulfilled this requirement. We excluded neurons that did not belong to distinctly separated clusters from further analysis. This procedure resulted in the identification of 17 clusters in the human fovea dataset (N = 481), 35 clusters in the human periphery dataset (N = 851), and 45 clusters in the macaque periphery dataset (N = 6228), i.e., we deliberately split the data into many smaller clusters. This procedure ensured that firing rate vectors reflecting noisy responses or light artifacts were grouped into their own clusters and firing rate vectors from different RGC cell types would not be erroneously grouped into the same cluster. After assessing each clusters’ mean responses to the light stimulus, we manually removed those that reflected noise or artifacts. This refinement resulted in 13 clusters for the human fovea (N = 347), 35 clusters for the human periphery (N = 851), and 42 clusters for the macaque periphery (N = 5844). We then reevaluated the trial-averaged and normalized firing rates of the neurons that remained after the initial analysis, applying UMAP once more, followed again by hierarchical clustering. In this second iteration, the hierarchical clustering algorithm was stopped at varying numbers of clusters (ranging between 2 and 40). To assess the quality of clustering, we calculated a silhouette score (*50*) for each potential number of clusters. The silhouette score, in conjunction with a visual inspection of the dendrogram — which visually depicts the distances between successive merges or fusions — outputted from the *AgglomerativeClustering* function, guided us on determining an appropriate cut-off for the number of clusters. This process resulted in 5 clusters for the human fovea, 10 clusters for the human periphery, and 7 clusters for the macaque periphery. Within our datasets, we identified On/Off parasol/midget cells characterized by their distinctive transient and sustained responses to increments/decrements of light, respectively (*33*). In the human fovea, the 5 clusters (N = 347) indicated On transient, On sustained, Off transient, Off sustained, and On sustained with elevated background activity (Fig. 2F). Transient responding cells were classified as parasol cells (Fig. 2F, cluster 4 and 5) and cells with sustained responses as midget cells (Fig. 2F, cluster 1-3). In the human periphery dataset (N = 746), we identified and merged clusters that displayed similar behavior to midget and parasol cells, ultimately yielding four distinct clusters: On midget, On parasol, Off parasol, On-Off cells (Fig. 2G, clusters 1-4). For the macaque periphery dataset (N = 4145), we selected the 4 clusters that best matched the response profiles of comparable cell types (Fig. 2H). The feature vectors of individual cells were finally plotted in a two-dimensional UMAP coordinate space, which was rotated to position the clusters corresponding to On cells at the top.

### Tracking propagating action potentials in recordings of spontaneous spiking activity

We reconstructed the average action potential (AP) waveform of each spike-sorted retinal ganglion cell (RGC) across the microelectrode array with the method described in ‘Recording spontaneous spiking activity with HD-MEAs’. The waveform of each neuron was represented by a 3-dimensional matrix W(x, y, t), where x and y were the electrode row and column of the HD-MEA, respectively. The center-to-center electrode distance was 17.5 μm. The time coordinate t was defined as the number of frames at 20 kHz resolution. We visualized W as a movie, where each pixel represented one electrode, and color represented the voltage value at this electrode. This visualization technique, (see supplementary video, Movie S1), enabled us to manually trace the AP’s trajectory within the video using a custom-built user interface in MATLAB. The exact moment of AP arrival at the different electrodes along its path was determined by identifying the mid-point between the minimum and maximum of the waveform at each location (Fig. 1C).

The result of this analysis was an AP trajectory, i.e., a set of space-time coordinates (x, y, t) that defined where the AP passed at what time. We excluded AP trajectories with less than three annotated space-time coordinates from further analysis. AP trajectories were smoothed and resampled utilizing Gaussian process regression (GPR, MATLAB function *fitrgp*, followed by *predict*, significance level, 0.05; prediction type, ‘*curve*’). This process was conducted separately for the *x* and *y* coordinates, yielding trajectories with space-time coordinates separated in time by Δt = 0.01 s. To estimate the speed of the AP propagation, we converted the two spatial coordinates (x, y) into a travel distance, d, by linearly integrating the distance between successive space-time coordinates along the AP trajectories. We then calculated the AP propagation speed for each RGC by a linear regression between the travel distance and travel time. As we noticed that the AP propagation speed was highly variable along the initial 200 μm closest to the soma, we excluded this initial part of the trajectory from the regression.

### Tracking propagating action potentials in recordings of light responses

The recordings of light evoked RGC responses necessitated a different recording strategy, because we wanted to spike sort - with high quality - a large number of RGCs simultaneously. Therefore, in this dataset, the average action potential waveforms could not be mapped over the entire HD-MEA but were constrained to a smaller area of the HD-MEA. We proceeded with the analysis of this data set as described in section ‘Tracking propagating action potentials in recordings of spontaneous spiking activity’ with the difference that the tracking was constrained to the smaller area. Neurons for which the axonal signal amplitude was insufficient for the tracking were removed from the analysis, but we did not exclude axons with tracked lengths below 200 μm per default. This resulted in the tracking of 113 midget cell axons (78 On and 35 Off), and 29 parasol cells axons (26 On and 3 Off) in the foveal data set. For the human periphery dataset, we tracked 102 On midget and 258 parasol cells (227 On and 31 Off) (Fig. 2I). For the macaque periphery dataset (Fig. 2J), we tracked 294 On midget cells and 224 On parasol cells.

### Analysis of axon trajectories and propagation speeds

We then grouped the reconstructed RGC axon trajectories based on the retinal location (quadrant and the distance from the optic disc) of the explants from which they originated. This process resulted in a total of 4758 tracked RGC axons: 1285 from human retinal explants containing the fovea centralis (10 donors; 11 explants, including foveola (N = 37), fovea (N = 1135), parafovea (N = 108), and perifovea (N = 5)), 1273 from human peripheral retinal explants (87 along the superior-inferior axis and 1186 along the naso-temporal axis; 9 donors and 20 explants), 128 from macaque retinal explants containing the fovea, and 2206 from macaque peripheral retinal explants (846 along the superior-inferior axis and 1354 along the naso-temporal axis; 11 specimens and 16 explants). The maximum length over which we could track AP trajectories were 1.67 mm for human fovea, 3.06 mm for human periphery, 1.96 mm for macaque fovea, and 3.33 mm for macaque periphery. For explants that contained the fovea centralis, we determined the position of the fovea centralis from the electrical activity recorded with the HD-MEA. To this end, we visualized the spiking activity of the explant as an image where each pixel represented an electrode and color coded the number of spikes detected at that electrode (Fig. S1). In these images, the ring-like structure of high RGC density around the umbo became clearly visible as a ring of high spiking activity, which allowed us to localize the position of the center of the fovea on the HD-MEA (Fig. S1, white triangle) for each foveal explant. We determined the direction of the optic disc by plotting all AP trajectories on top of each other and observing the characteristic bending pattern. We then rotated and shifted all AP trajectories so that the center of the fovea was at the origin and the optic disc in the direction of 0°. This procedure effectively registered all AP trajectories from different explants containing the fovea centralis in a shared coordinate system. We quantified the relationship between the AP propagation speeds and the positions of the corresponding RGC somas in relation to the fovea. To this end, we defined the ‘RGC angular position’ as the angle formed by two lines: one connecting the location of the first space-time coordinate of the axonal trajectory with the position of the fovea centralis, and the other extending from the fovea centralis to the optic disc (i.e., fovea-optic disc axis, defined as 0°). We grouped the RGC angular position into bins of 30°. Within these bins, we computed the mean and standard error of the mean (SEM) of the AP propagation speeds, as illustrated in Fig. 1G.

### Analysis of AP propagation speed distributions

We constructed AP propagation speed distributions of RGC axons as histograms with 50 bins, as depicted in Fig. 2A and Fig. 2C, normalizing them to probability density functions. The distributions revealed a bimodal pattern, suggesting the presence of at least two distinct RGC populations. To deconvolve these populations, we fitted a Gaussian mixture model (utilizing MATLAB’s *fitgmdist* function) with two components (k = 2, 1000 optimization iterations) independently to the speed distributions of each retinal region (center, mid- and far-periphery).

### Electron microscopy sample preparation and imaging

Fixed human retinal sections (4% PFA) were rinsed once in Cacodylate buffer (0.1 M, pH 7.3) for 10 min. After two additional washes in Cacodylate buffer, sections were post-fixed in 1% osmium tetroxide, 0.8% potassium ferrocyanide in 0.1 M Cacodylate buffer for 1 hour at 4 °C. Sections were then rinsed several times in Cacodylate buffer and ultrapure distilled water then, *en bloc* stained with 1% aqueous uranyl acetate for 1hr at 4 °C in the dark. After several wash steps in ultrapure distilled water, sections were dehydrated in an ethanol series (30, 50, 75, 96, and 100%) at 4 °C followed by three additional washes of absolute ethanol. Sections were then washed in acetone and finally embedded in a mixture of resin/acetone first and then in pure Epon 812 resin (Embed 812-EMS) overnight. Sections were flat embedded first using Adhesive frames (Gene Frame, 25 μL; Thermo Fisher Scientific). Polymerization was carried out for 48 h at 60 °C. Each polymerized section was then cut into a small strip. Each strip was re-embedded in Epon resin and polymerized for an additional 2 days at 60 °C. The position of the sample within the embedding was based on the orientation of the axons within the sample. We positioned the sample so that the axons were cut in cross-section. 70 nm ultra-thin sections were obtained with a diamond knife and collected on copper slot grids, coated with Formvar film and carbon layer, stained with uranyl acetate and lead citrate, and observed into a Talos L120C G2 (Thermo Fisher Scientific™) operated at 120 kV, equipped with a 4k x 4k Ceta CMOS camera. The SerialEM (*51*) program was used for automated image acquisition of a large area from serial sections (polygon). Polygons were all acquired at the magnification of 8500x. It is important to mention that resin-embedding can lead to the shrinkage in biological samples, potentially impacting the estimation of axonal diameters compared to those obtained in vivo or through alternative methods (*52*). However, for the purpose of this study, the relative difference of axonal diameters was the relevant quantity, not the absolute diameters.

### Estimation of axon diameters in electron microscopy images

Large polygonal areas, acquired via transmission electron microscopy (TEM), were cropped into 3000 x 3000-pixel images using ImageJ software (Fiji distribution). Utilizing Cellpose 2.0, we trained a custom segmentation model on a random subset of these images (*53*). Subsequently, each TEM image was processed using this custom-trained model. Remaining errors in the output of the automatic segmentation procedure were corrected through manual curation. The segmented outlines from each image were then exported and analyzed with ImageJ (Fiji). In Fiji, we fitted the Cellpose-generated outlines with ellipses, and used the lengths of the minor axes of these ellipses as the axon diameters. This was done to ensure that a tilt of an axon with respect to the imaging plane, which would elongate the outline of the axon in the direction of the tilt, would not result in a bias towards larger axon diameters.

### Assessment of retinal nerve fiber layer thickness via optical coherence tomography

To assess the thickness of the retinal nerve fiber layer (RNFL, Fig. S2B), we conducted optical coherence tomography (OCT) imaging using a Zeiss Cirrus HD-OCT machine. The Optic Disc Cube 200x200 scan protocol was employed for imaging. RNFL thickness measurements were obtained using the device’s built-in segmentation algorithm.

### Psychophysics: human foveolar reaction time

To ensure spatially resolved retinal photo stimulation for simple reaction time (RT) measurements in humans, a custom-built adaptive optics scanning laser ophthalmoscope (AOSLO) was employed. In an AOSLO, the retina and stimulus location can be resolved with sub-cellular resolution for precise photoreceptor-targeted psychophysical examination. Technical details of the AOSLO instrument and stimulation techniques have been described previously (*54*). In short, carefully controlled doses of 543 nm light were briefly flashed against the 840 nm, 0.85-deg field of view raster of the AOSLO to hit either only a single cone photoreceptor, or a small group of cones. The light distribution on the retina in the small stimulus was 1.8 µm full width at half maximum (i.e., FWHM) considering 0.03 Diopter of residual defocus, and 9.2 µm FWHM in the larger stimulus. Stimulus duration was 125 µs for the small, and 1,126 µs for the large stimulus. Stimulus duration also dictated the total amount of stimulus power delivered to the retina, which was 0.3 nW for the small, and 12 nW for the larger stimulus. In each trial, a stimulus would appear randomly displaced in a central subfield of the imaging raster, producing a close to normally distributed stimulus delivery location relative to the foveal center. The stimulus delivery location was corrected for transversal chromatic offsets (*55*).

To avoid any adaptation or anticipation of the next stimulus delivery, a variable time interval of 0.5 to 1.5 seconds was added after the trial onset, initiated by a keyboard press of the participant. RTs were measured as the time between stimulus delivery onset, detected in the drive signal to the acousto-optic modulator by the trigger function of a fast oscilloscope (MSO-X 3054A, Agilent Technologies), and a detection response. Participants indicated stimulus detection via a button press of a custom-made hardware microswitch. Millisecond resolution without temporal interference was achieved by using an Arduino microcontroller (Arduino AG, Italy), measuring the delay between the stimulus onset indicated by the oscilloscope trigger and the participant’s button press. The measured RT served then as input to a second computer running the AOSLO experiment via a MATLAB interface and saved to a log-file.

Foveal RTs were measured in one female and two males (ages: 33, 34 and 44) with no known retinal conditions. Mydriasis and cycloplegia were induced by instilling one drop of Tropicamide into the lower eyelid 15 minutes prior to experimentation and subsequent re-dropping if necessary.

Individual stimulus positions were recovered from single AOSLO image frames and registered to a high signal-to-noise ratio average image of each of the participant’s foveolar center to ensure a precise retinal stimulus localization. The high-quality retinal images were derived by spatially registering and normalizing about 150 individual AOSLO image frames by strip-wise image registration (*56*). In these images, the location of each cone was semi-manually annotated to compute a 2-dimensional map of cone density. The center of the fovea (i.e., zero eccentricity), was defined as the location of the cone density centroid (CDC), which was computed as the weighted center of the 80% density isoline contour of the full density map (*30*). Out of a total of 3000 trials, 348 (11.6%) had to be discarded because the foveolar image could not be registered to the foveolar center, resulting in uncertain retinal stimulus locations. An additional 288 trials (10.9% of the remaining 2652) were removed containing implausible RTs shorter than 140 ms, probably being a product of stimulus anticipation. In total 636 trials (21.2% of 3000) were excluded from the analysis, leaving 2364 valid trials. All psychophysical and imaging procedures were conducted with approval of the independent Ethics committee of the Medical Faculty of the Rheinische Friedrich-Wilhelms-University Bonn (Lfd-Nr. 294/17) and adhered to the tenets of the Declaration of Helsinki.

### Reaction times: temporal vs nasal

To enhance the statistical power of our analysis on reaction times (RTs) measured using adaptive optics scanning laser ophthalmoscopy (AOSLO) across three participants, we initially identified and removed outlier trials (*rmoutliers*, MATLAB) constituting 0.6% of all 2253 trials.

Following outlier removal, we normalized the data for each participant. To normalize each participant’s data (both nasal and temporal), for each participant we subtracted the mean RT over all trials. We then computed each participant’s pooled standard deviation: We assigned the trials of each participant to two distinct regions (temporal and nasal) based on each trial’s position relative to the cone density centroid (CDC) of the respective participant. We computed the standard deviations (STDs) of the RTs for the nasal and temporal region separately, yielding two STDs per participant. For each participant, we then computed their pooled standard deviation as follows:

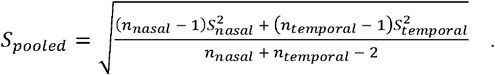

To normalize the RTs and we then divided each participants RTs by their respective pooled standard deviation. Following normalization, the data from all participants was aggregated into a single dataset keeping the original assignments into temporal and nasal regions of each participant. We then conducted a power analysis using MATLAB’s *sampsizepwr* function for a two-sample t-test to ascertain the smallest increase in RT of the temporal group that could be reliably detected given the number of trials. Because the two regions could have different number of trials, we used the smaller of the two trial numbers. The power analysis was performed at a fixed significance level (alpha) of 0.05. We then calculated the smallest effect size (temporal RTs larger than nasal RTs), which yielded a power (beta) of at least 0.8. The power analysis was performed independently for the two different light stimulus sizes. The analysis revealed that an effect size of 7.5 ms (small stimulus, N = 517) and 5.5 ms (large stimulus, N = 489) corresponded to the given significance and power levels.

### Estimating orientations of axon bundles from micrographs of human whole mounts

The reconstructed image of the human whole-mount retina had a resolution of 219 pixel per mm and a size of 12k x 12k pixels, which encoded contrast in values ranging from 0 to 255. We determined the local orientation of axon bundles in a window of 201 x 201 pixels, which we moved in steps of 50 pixels over the image. We set small contrast values below 20 to a value of 0 to remove noise on the black background of the image and ignored windows with a median contrast value below 20. In each window, we used the method detailed in (*57*) to determine the bundle orientation. Briefly, the method calculated the 2D Fourier transformation of the image within the window, which decomposed the image into a set of 2D sine waves, characterized by direction, spatial phase, and amplitude. Low spatial frequency components usually reflect the background of the image within the window and other unwanted image features like uneven illumination. High-frequency components are often dominated by noise. The method, therefore, relied on a spatial bandpass filter to block those components. After filtering, the method yielded the orientation of the spatial frequency components with the highest amplitude. The orientation corresponding to this frequency component was then returned as the orientation of the axon bundles.

### Model of axonal trajectories in the human retinal nerve fiber layer

We modeled the geometry of the human retina as a sphere of 12 mm radius and defined a point on this sphere as the origin in polar coordinates. Opposite to the origin, we removed a spherical cap from the sphere, so that the ora serrata, i.e., the location of the cut, was at a geodesic distance of 125° (or 26.18 mm arclength) from the origin. In our geometry, the fovea was located at (13 mm, -4 mm) and the optic disc at (-10 mm, 0 mm). At the location of the optic disc and fovea we removed a spherical cap (i.e., inserted a hole) in the eye of 0.6 mm and 0.2 mm radius, respectively. For numerical reasons we found it easier to work with a three-dimensional geometry and, therefore, gave the retina a thickness of 0.24 mm. This procedure defined a geometry of a spherical shell with three circular holes representing the anterior segment of the eye, the fovea, and the optic disc. We constructed a 3-dimensional mesh of the resulting geometry using MATLAB’s PDE toolbox. On this mesh, we solved equation (1).

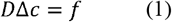

The parameter *D* represented the diffusivity of the retina, *c* was the unknown concentration of the substance that guides axonal growth, and *f* was a function which implemented the spatial extent of the source at the fovea defined as follows

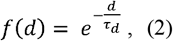

where *d* represented the distance from the fovea, and *τ*_*d*_ was a parameter that controlled how fast the strength of the source at the fovea decayed with distance from its center. Sinks and sources were furthermore defined via Dirichlet boundary conditions at the borders of the three holes: (i) for the ora serrata, *b*_*OS*_, (ii) for the fovea *b*_*F*_, and (iii) for the optic disc *b*_*OD*_. This resulted in a model with 5 parameters (*D, τ*_*d*_, *b*_*OS*_, *b*_*F*_, *b*_*OD*_). The model was solved using MATLAB’s *solvepde* function. The solution defines the concentration *c* across the retina. The directional component of the spatial gradient of *c* defined the axonal directions. To calculate axonal trajectories, we used MATLAB’s *stream2* function that received the axonal directions as input.

### Fitting the model to axonal orientations

We fitted the model parameters to the regions of whole-mount immunolabeled images in which we manually annotated the location of the optic disc and fovea. We then shifted, rotated, and scaled the model RNFL, so that the model fovea and the model optic disc were coinciding with those visible in the whole mount image. In contrast to the model, which described the local directions of the axons in the range of 0° to 360°, the axonal orientations calculated from the whole-mount images were defined in the range of 0° to 180°. To make these two quantities (direction and orientation) comparable, we converted the model directions to orientations. This was achieved by subtracting 180° from all directions between 180° and 360°. We then fitted the model to the axonal orientation within the foveal region by minimizing the average circular distance between modelled orientation and the local orientations of the axon bundles, extracted from the image of the whole mount. We applied this procedure to two whole-mount immunolabeled human retinae. The fitted values for the parameters were *D* = 0.022, *τ*_*d*_ = 1.74, *b*_*OS*_ = 945.66, *b*_*F*_ = 929.77, *b*_*OD*_ = −3.90 for the first whole-mount, and, *D* = 0.026, *τ*_*d*_ = 1.82, *b*_*OS*_ = 854.37, *b*_*F*_ = 839.97, *b*_*OD*_ = −4.10, for the second whole-mount. The resulting *R*^2^ values of the fits amounted to 0.91 and 0.95, respectively. However, when we used the parameter values of the fit to the first whole-mount to model the axon trajectories of the second whole-mount, the resulting *R*^2^ value was still 0.95 emphasizing how similar the fitted parameter values were for the two different human retinae.

### Modelling RGC axonal density across the retina

The model specified the pathways of RGC axons from their soma of origin to the optic disc. It did not specify how many RGC somas were present at each retinal location. We modeled the RGC soma density using the following function:

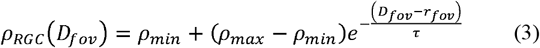

where *ρ*_*RGC*_ was the RGC density at a distance *D*_*fov*_ from the center of the fovea, *ρ*_*min*_ and *ρ*_*max*_ were the minimal and maximal RGC densities across the retina, *τ* was a spatial scaling parameter and *r*_*fov*_ was the radius of the umbo, the area in the center of the fovea devoid of RGCs. To reflect the human RGC density described in (*58, 59*), we set the parameters to 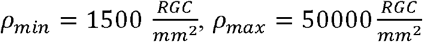, *τ* = 1.43 and *r*_*fov*_ = 0.2 *mm* We then applied the Delaunay triangulation to overlay the model retina with 167,000 triangles of approximately equal size. We determined the count of RGC somas within each triangle by integrating equation (3) across the triangle’s surface. Subsequently, the model enabled us to estimate, for each triangle, the axonal pathway connecting the triangle’s centroid to the optic disc. Thus, each of the 167,000 axonal pathways was linked to a certain number of RGC axons, which followed this pathway closely. To ascertain the quantity of axons traversing between two closely spaced points on the retina, designated as A and B, we identified the field lines crossing the line connecting A and B and summed up the corresponding RGC axon counts.

---

**Supplementary Fig. 1:**
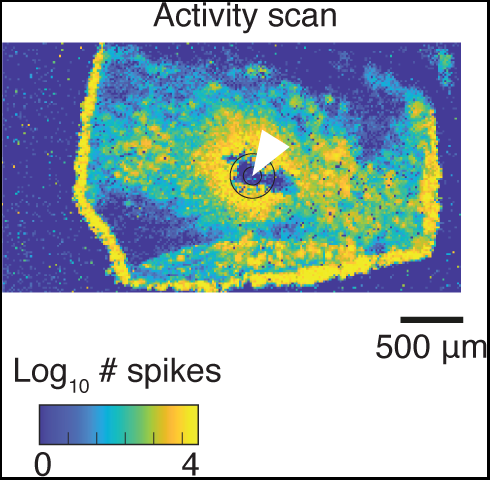
Example of electrical recording of ex vivo human foveal retinal explant. Electrical activity of an example of ex vivo human foveal explant. Color encodes the spike rate recorded at each electrode. The electrode position is encoded by the pixel position. White arrow: Center of the fovea. Concentric black circles: Umbo (small) and foveola (large).

**Supplementary Fig. 2:**
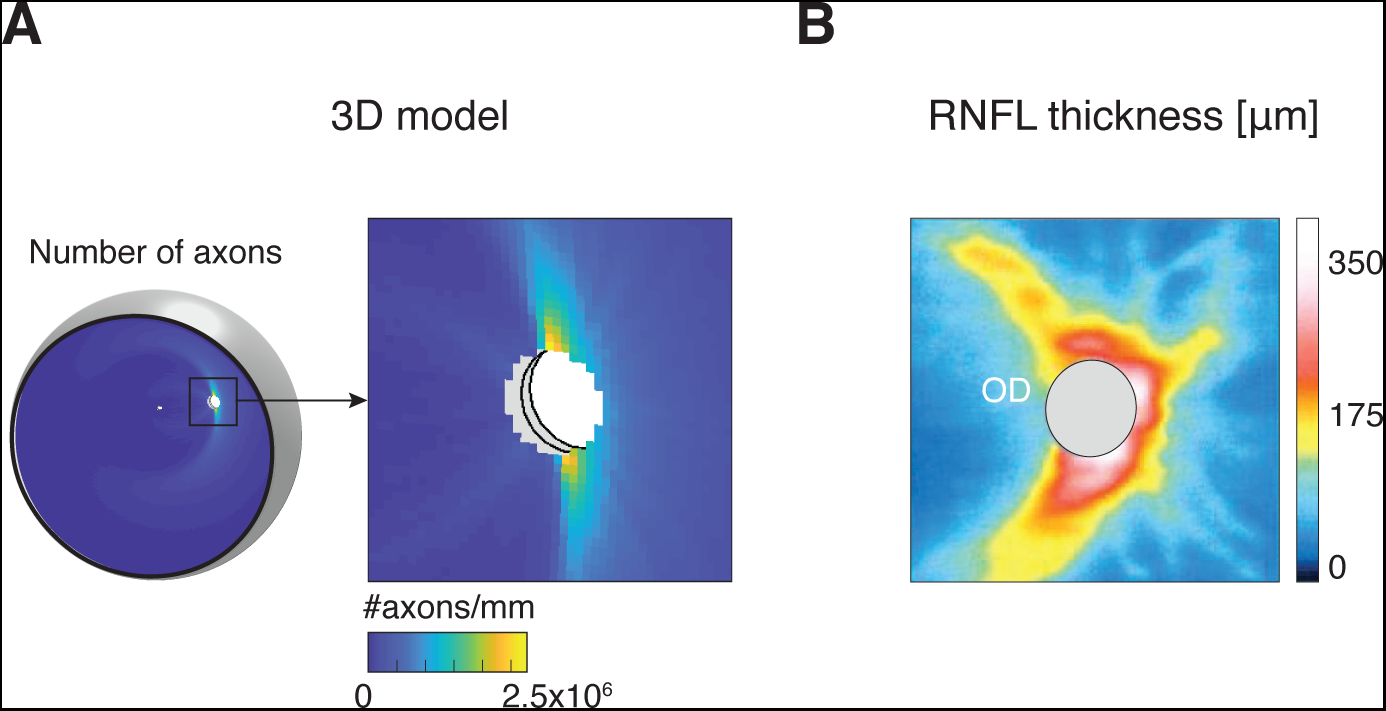
Retinal nerve fiber layer thickness. (**A**) Number of axons at each location within the retina. Same as shown in Fig. 3D. Right: Zoom on the region containing the optic disc (white hole in the center). (**B**) Thickness of the RNFL of one of the authors measured with OCT. Grey ellipsoid: Optic disc.

**Supplementary Movie 1: Propagation of a single foveal RGC action potential across the HD-MEA**.

Average voltage recording of an action potential of a foveal RGC reconstructed from action potentials recorded with different electrode configurations. Data was bandpass filtered and processed to optimize visualization. Color encodes voltage at each electrode (arbitrary unit). The electrode position is encoded by the pixel position.

